# fMRI Correlates of Reaction Time Prolongation during intentional False Responding; an inter-individual difference study

**DOI:** 10.1101/089847

**Authors:** Morteza Pishnamazi, Maral Yeganeh Doost, Habib Ganjgahi, Hamed Ekhtiari, Mohammad Ali Oghabian

## Abstract

Reaction time (RT) is chiefly longer when people lie. However, the baseline speed in answering questions and the amount of RT prolongation during lying show considerable amount of inter-individual variability. In the current study, we exploited this fact to glean insights on the contribution of each lie-related brain region to hampering of response speeds when people try to be deceitful. In an event-related fMRI session, participants were interrogated by yes-no autobiographical questions and were instructed to intentionally provide false responses to a pre-selected subset of questions. Data from twenty healthy volunteers were analyzed. *Baseline speed* [RT_truth_] and *relative appended lie RT* [(RT_lie_ − RT_truth_) ⁄ RT_truth_] measures were calculated for each participant and were included in the group level analysis of [lie > truth] BOLD contrasts. Lying RTs were significantly longer than truth telling RTs. Lie-related increase in activity of right ventrolateral prefrontal cortex (VLPFC) and bilateral paracingulate cortex correlated with the baseline speed of participants, while the increase in activity of Left VLPFC, left lateral occipital cortex and bilateral anterior cingulate areas directly correlated with the amount of lying reaction time cost. Activity within bilateral posterior cingulate cortex and right insular cortex inversely correlated with lying RT-cost. Bilateral supplementary motor areas, internal capsule white matter and left angular gyrus showed lie-related increase in activity but did not correlate with either of behavioral measures. Provisional implications regarding the contribution of these regions to RT prolongation and their cognitive role in deceitful behavior are discussed.

## 1. INTRODUCTION

With the advent of functional magnetic resonance imaging (fMRI), there has been a fast growth in the neuroimaging literature of deception. Studies commonly aimed to reveal the neural correlates of deception by contrasting brain activities recorded under conditions of instructed lying versus conditions of truth telling (Ganis, Kosslyn, Stose, Thompson, & Yurgelun-Todd, 2003; Langleben et al., 2002; T. M. C. Lee et al., 2002; Spence et al., 2001). Early experiments were followed by series of studies that tried to reinforce previous findings by using life-like task designs, such as mock crime scenarios (Kozel et al., 2005, 2009) and deceptive games (Sip et al., 2010, 2012). Results have been comparatively consistent, showing that areas in ventrolateral prefrontal cortex (VLPFC), dorsolateral prefrontal cortex (DLPFC), insular cortex, inferior parietal lobule (IPL) and anterior cingulate cortex (ACC) are more active during lying and deception (Abe, 2011; Christ, Van Essen, Watson, Brubaker, & McDermott, 2009; Farah, Hutchinson, Phelps, & Wagner, 2014). However, not enough care has been put into interpreting the specific function of each region; nor has been there enough consideration of the nuisance variables that could confound the fMRI comparisons between lie and truth conditions (Sip, Roepstorff, McGregor, & Frith, 2008).

Providing false responses is a complex cognitive task that involves processes additional to those used when telling the truth (Williams, Bott, Patrick, & Lewis, 2013) and demands higher mental effort (Caso, Gnisci, Vrij, & Mann, 2005; Vrij, Granhag, Mann, & Leal, 2011). To formulate a false response, one requires to first activate the truth and then modify it (Debey, De Houwer, & Verschuere, 2014). This adds the steps of response inhibition, task switching and response planning (Debey, Liefooghe, De Houwer, & Verschuere, 2014; Gombos, 2006; Walczyk, Roper, Seemann, & Humphrey, 2003). Besides, in comparison with truth telling, lying depends more heavily on working memory and maintained attention (Gombos, 2006; Vendemia, Buzan, & Simon-Dack, 2005). These higher cognitive demands is reflected in the longer reaction times (RTs) associated with deceptive responses (Verschuere, Suchotzki, & Debey, 2015). Repeated studies show that RT is chiefly longer when people lie (Marston, 1920; Sheridan & Flowers, 2010; Vendemia et al., 2005; Walczyk et al., 2003). However, the baseline speed in answering questions and the amount of RT increment imposed by the act of lying (*‘RT-cost’*) differ from one individual to another. In the study by Farrow and colleagues (Farrow, Hopwood, Parks, Hunter, & Spence, 2011) subjects with higher memory ability had lower absolute truth RTs but the RT difference scores (lie RT minus truth RT) were adversely affected, showing a positive correlation between memory ability and the RT-cost of lying. Visu-Petra and colleagues (Visu-Petra, Miclea, Buş, & Visu-Petra, 2014; Visu-Petra, Miclea, & Visu-Petra, 2012) studied the relation between inter-individual differences in executive functions (inhibition, shifting, working memory) and the latency of deceptive responses. Subjects with better inhibitory skills had faster absolute lie RTs but RT difference scores showed no correlation with any executive function measure. Despite the intuitive involvement of arousal and emotion mechanism in deception, behavioral experiments report mixed results about the association between deceptive RTs and measures of anxiety (Visu-Petra et al., 2012), personality (Verschuere & in ´t Hout, 2016; Visu-Petra et al., 2014) or motivation (Kleinberg & Verschuere, 2016; Varga, Visu-Petra, Miclea, & Visu-Petra, 2015).

Superior executive skills seem to be linked with faster baseline speed in answering questions and lower RT-cost of lying. This association is supported by neuroimaging findings of higher activity within multiple executive function-related regions in frontal cortex during lying (Christ et al., 2009). However, the exact relationship between activity within each area and reaction times is not clear. In the current study we aspire to exploit the inter-individual variance in reaction times to glean insight on the neural mechanisms underlying prolongation of reaction times when people lie. To that end, we enrolled 25 healthy volunteers in an fMRI experiment. We recorded reaction times and blood oxygenation level dependent (BOLD) brain activations while subjects were interrogated by a set of yes-no questions and intentionally provided false responses to a preselected subset of them. Based on reaction time recordings we calculated measures representing each subjects’ baseline speed and RT-cost of lying. We investigated the correlation of RT-measures with the amount of BOLD activation difference between lying and truthful responding conditions across subjects. The critical question of interest is to specify which brain regions undertake cognitive processes exclusive to lying (e.g. response inhibition, task switching). Activity of such a region is expected to only correlate with lie RTs. On the other hand, activity in regions commonly employed by both truthful and false responding (e.g. attention, working memory) is expected to correlate with truth RTs and lie RTs similarly.

## 2. MATERIALS AND METHODS

### 2.1. Participants

Twenty-five healthy, right-handed male volunteers (age 21-30) were recruited and provided informed consent. All participants went through a standardized medical interview. Exclusion criteria were any history of psychiatric or neurological disorder, use of any medications during last week, and general MR safety contraindications. Five participants were excluded from data analyses (three failed to perform experimental procedures adequately; two because of technical problems in data gathering). The ethical committee of Tehran University of Medical Sciences approved all procedures.

### 2.2. Procedure

In resemblance to the lying paradigm used in the study by Nuñez and colleagues (Nuñez, Casey, Egner, Hare, & Hirsch, 2005), our task consisted of yes-no autobiographical questions (e.g. “Do you own a car?”) and required intentional false responding. First, participants provided truthful yes-no answers to 20 autobiographical questions. We asked subjects to freely choose half of questions. Next, we instructed them to lie about these pre-selected questions for the rest of the experiment. Prior to main fMRI session, a 5-minute training was run outside the scanner to ensure participants’ familiarity with task procedure. Total duration of main fMRI session was 16 minutes. We employed event-related task design. Each of 20 questions was presented 5 times in counterbalanced random order. Each question was presented for 2 seconds, followed by a jittered inter-stimulus interval ranging from 3.5 to 11.5 seconds during which a central fixation sign was displayed. Participants’ responses and reaction times were recorded. Three types of event could happen: ‘truthful’ answer to questions, intentional ‘false’ answers, and ‘mistakes’ where subjects failed to provide appropriate response based on their template. Event types were determined post-hoc based on each subjects’ original responses to questions and their pre-selected lying subset.

### 2.3. fMRI Data Acquisition

Images were acquired using 3.0 T Siemens Magnetom Tim Trio full-body scanner with 12-channel head coil. Functional T2*-weighted images were collected using gradient echo-planar imaging (TR = 3000ms, TE = 30ms, flip angle = 90°, FOV = 192 mm, matrix = 64 × 64, voxel size = 3 × 3 × 3 mm). 40 contiguous axial slices provided whole-brain coverage. Three additional images were included at the start of each run to allow signal stabilization and were excluded from analysis. Prior to the functional scan, T1-Weighted high-resolution structural image was acquired (TR = 1800ms, TE = 3.44ms, flip angle = 7°, FOV = 256 mm, matrix = 256 × 256, voxel size = 1 × 1 × 1 mm). This image was used for anatomical coregistration and normalization.

### 2.4. Reaction Time Analysis

Reaction times of each subject were averaged over truthful events (RT_truth_) and false responding events (RT_lie_). Mistake events and outlier RTs (values that deviated more than 1.5 inter-quartile ranges from the upper and lower quartiles of each subject) were excluded from these calculations. We aimed to investigate the relationship between the amount of RT prolongation during lying and the amount of activity in lie-related brain regions. We employed average RT_truth_ of each subject as the measure of baseline speed. Inter-individual variance in RT_truth_ values reflects differences in general dexterity of subjects in performing task requirements. Competence at cognitive skills specific to the act of deception is not expected to affect the baseline speed in answering questions. As the measure of RT-cost of lying we calculated each subjects’ *‘relative appended lie reaction time’* [(RT_lie_ − RT_truth_) ⁄ RT_truth_]. Expressing lying RT-cost as a fraction of baseline speed nulls the effect of dexterity and accentuates the influence of deception-specific cognitive processes in determining RT_lie_ of an individual.

### 2.5. fMRI Data Analysis

FMRI data processing was carried out using FEAT Version 5.98, part of FSL (FMRIB’s Software Library, www.fmrib.ox.ac.uk/fsl) (Jenkinson, Bannister, Brady, & Smith, 2002). Preprocessing steps consisted of: brain extraction (Smith, 2002), motion correction (Jenkinson et al., 2002), slice-time correction, Gaussian spatial smoothing (FWHM= 5 mm), and high-pass temporal filtering of the time series. Voxel time series were modeled by general linear model. Regressors for each event type (‘truthful’, ‘false’, ‘mistake’) were convolved with canonical hemodynamic response function. BOLD contrast of [lie > truth] was calculated for each subject. Parameter estimates from single subjects were entered in random-effect group analysis (Woolrich, Behrens, Beckmann, Jenkinson, & Smith, 2004). *Baseline speed* measures [RT_truth_] and *lying RT-cost* measures [(RT_lie_ − RT_truth_) ⁄ RT_truth_] were included in the group-level analysis as between-subject regressors. This allowed us to perform voxel-wise whole-brain search for voxels where the BOLD activity difference between lie and truth events correlated with the inter-individual variability in baseline speed and RT-cost. For statistical inference, Z statistic images were thresholded at *Z* > 2.3; corrected cluster-significances of *p* < 0.05 were deemed meaningful.

## 3. RESULTS

### 3.1. Behavioral results

Frequency of mistakes was 10.2% on average (*SD* = 9.03), indicating that participants adequately adhered to their template and task instructions. RT_truth_ was 1.81 seconds on average (*SD* = 0.52) while mean RT_lie_ was 1.94 seconds (*SD* = 0.61). This difference was statistically significant based on paired-samples t-test [*t*(19) = 2.75, *p* = 0.013, effect size = 0.6 cohen’s *d*]. As expected, there was considerable amount of inter-individual variability among participants both in the baseline speed of answering truthfully (range: 1.12 – 3.18 seconds; Figure 1.a) and the difference between RT_lie_ and RT_truth_ (range: -0.24 – 0.63 seconds; Figure 1.b). *Relative appended lie RT* [(RT_lie_ − RT_truth_) ⁄ RT_truth_] ranged from -0.11 to 0.31.

**Figure 1.**
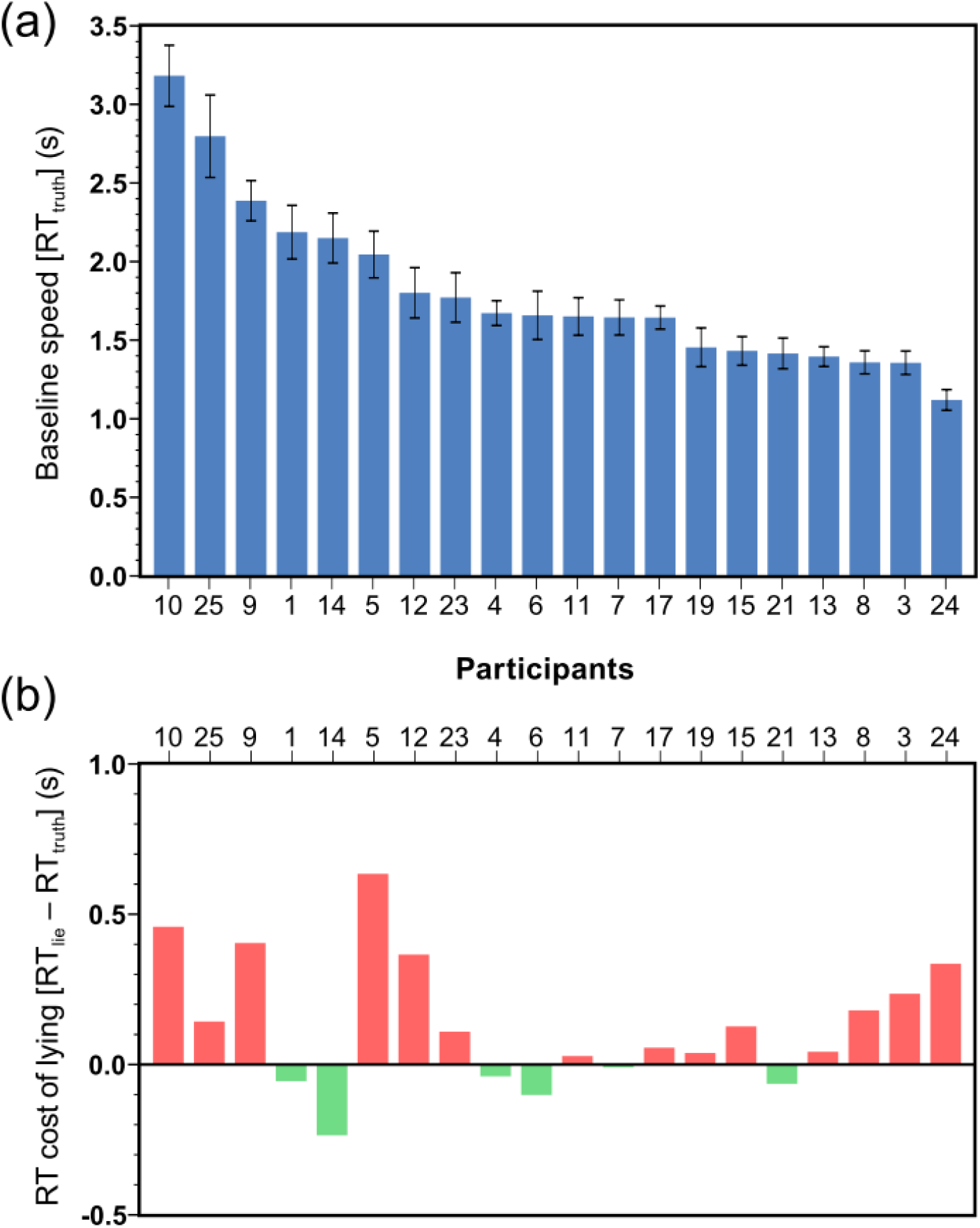
Participants showed considerable inter-individual variability both in the baseline speed and in the RT-cost of lying. **(a)** Average reaction time (RT) in truth trials, sorted in descending order. This value was used as the baseline speed measure of each participant in the fMRI analysis. Error bars show ±2 standard error of the mean. **(b)** Difference between mean RT in lie and truth trials [RT_lie_ − RT_truth_]. This value was divided by the RT_truth_ of each participant to yield *relative appended lie RT*, which was used as the lying RT-cost measure of each participant in the fMRI analysis. Bars colored green denote subjects with faster RT in lie trials.

Correlation between RT_lie_ and RT_truth_ of participants was highly significant [*r* = 0.941, *p* < 0.001]. On the other hand, the *relative appended lie RT* and RT_truth_ values were not correlated [*r* = 0.005, *p* = 0.984]; this allowed us to independently estimate their correlation with lie-related brain activations (Mumford, Poline, & Poldrack, 2015).

### 3.2. Functional imaging results

Table 1 presents the results of group-level contrast of [lie > truth] BOLD parameter estimates. We classified this set of anatomical areas into distinct subsets based on how their BOLD signal change correlated with behavioral RT-measures (Table 1, Figure 2). Activity in right inferior frontal gyrus (IFG) (corresponding to right VLPFC) and bilateral paracingulate cortex showed positive correlation with baseline speed measure [RT_truth_]. Left IFG (left VLPFC), left lateral occipital cortex (LOC), and bilateral anterior cingulate cortex (ACC) exhibited positive correlation with *relative appended lie RT measure* [(RT_lie_ − RT_truth_) ⁄ RT_truth_]. Areas showing negative correlation with this measure were bilateral posterior cingulate (PCC) and right insular cortex (Figure 2.a). A subset of areas showed higher BOLD activity during lying but the amount of their BOLD signal-change did not correlate with either of behavioral RT-measures. These areas were bilateral supplementary motor area (SMA), internal capsule white matter (ICWM), and left angular gyrus (AG; in the posterior segment of inferior parietal lobule, IPL) (Figure 2.b).

**Table 1.** Brain regions showing [lie > truth] BOLD effect, classified based on the correlation of their activity with *baseline speed^a^* and *lying RT-cost^b^* measures

|  | Brain region | H | Cluster size<br>(voxels) | Peak Z-value | Peak Coordinates<br>(MNI) |  |  | *p* value<sup>c</sup> |
| --- | --- | --- | --- | --- | --- | --- | --- | --- |
|  |  |  |  |  | x | y | z |  |
| **Baseline speed, Positive correlation** | Paracingulate gyrus | L & R | 1072 | 4.07 | -2 | 26 | 42 | < 0.001 |
|  | Middle frontal gyrus | L | 547 | 3.7 | -36 | -2 | 56 | 0.008 |
|  | Inferior frontal gyrus (VLPFC) | R | 515 | 3.38 | 42 | 24 | 20 | 0.012 |
| **Lying RT-cost, Positive correlation** | Inferior frontal gyrus (VLPFC) | L | 1520 | 4.4 | -54 | 20 | 24 | < 0.001 |
|  | lateral occipital cortex | L | 916 | 3.73 | -34 | -60 | 60 | < 0.001 |
|  | Anterior cingulate cortex | L & R | 855 | 4.01 | 0 | 20 | 36 | < 0.001 |
| **Lying RT-cost, Negative correlation** | Posterior cingulate cortex | L & R | 858 | 3.55 | 8 | -10 | 42 | < 0.001 |
|  | Insular cortex | R | 768 | 3.87 | 36 | 4 | 10 | 0.001 |
|  | Post central gyrus | R | 688 | 3.55 | 54 | -18 | 44 | 0.001 |
|  | Cerebellum nucleus V | R | 517 | 3.21 | 30 | -34 | -36 | 0.01 |
|  | Posterior middle temporal gyrus | R | 446 | 3.17 | 60 | -48 | 2 | 0.023 |
| **No correlation** | Internal capsule white matter | L & R | 1249 | 4.01 | 2 | -6 | 4 | < 0.001 |
|  | Supplementary motor area | L & R | 519 | 3.5 | -4 | 4 | 70 | 0.011 |
|  | Angular gyrus (IPL) | L | 448 | 3.22 | -54 | -56 | 38 | 0.026 |

<sup>a</sup> Average speed in answering questions truthfully [RT<sub>truth</sub>] was used as representative of participants' baseline speed.
<sup>b</sup> *Relative appended lie RT* [(RT<sub>lie</sub> – RT<sub>truth</sub>) ÷ RT<sub>truth</sub>] was used as representative of RT-cost of lying.
<sup>c</sup> Z statistic images were thresholded using clusters determined by $Z > 2.3$ and a corrected cluster significance of $p < 0.05$ .
H, hemisphere; IPL, inferior parietal lobule L, left; MNI, Montreal neurological institute; R, right; RT, reaction time; VLPFC, ventrolateral prefrontal cortex.

**Figure 2.**
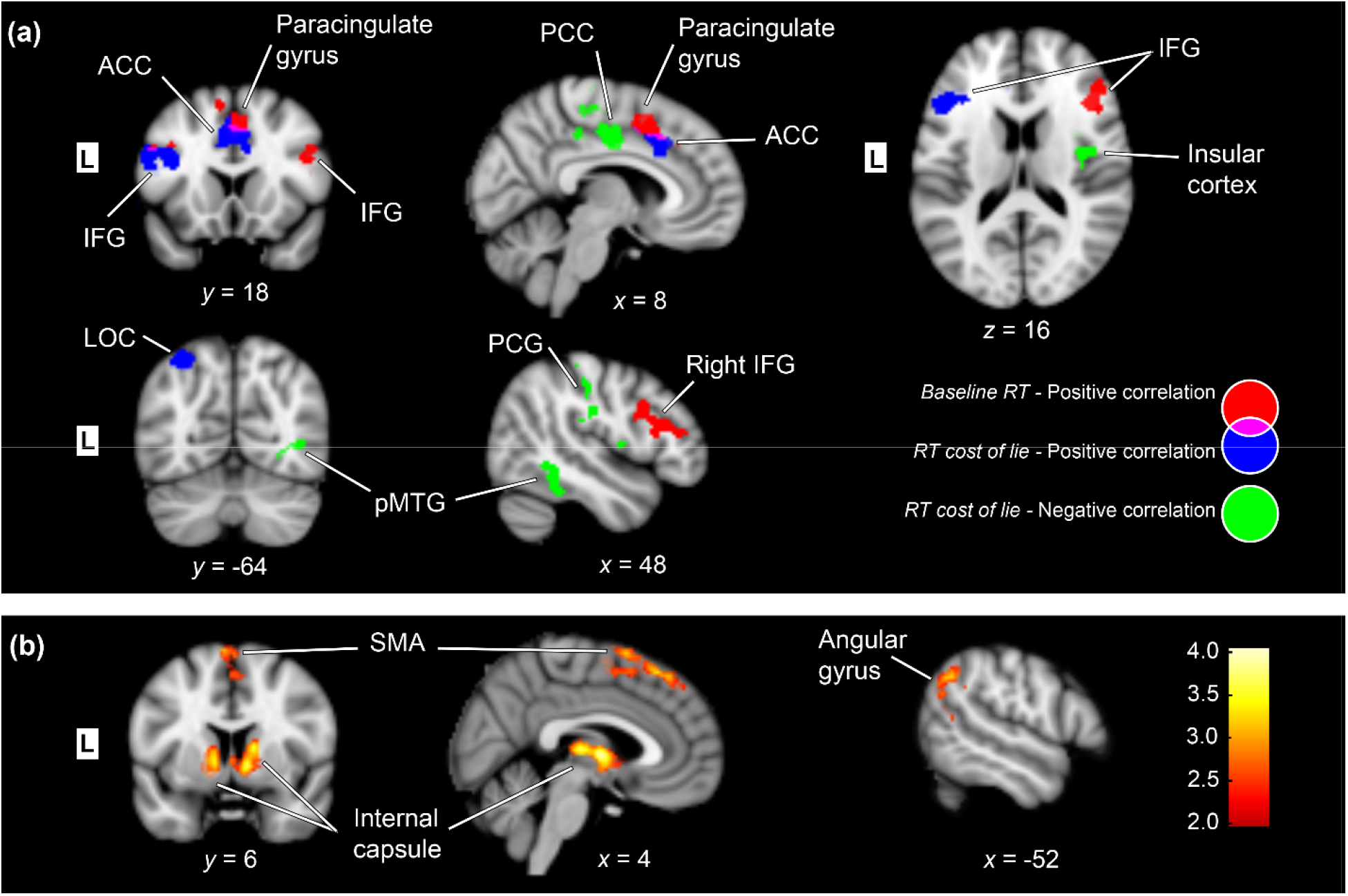
Group-level results of [lie > truth] BOLD contrast and correlations 487 with behavioral reaction time (RT) measures. Z statistic images were thresholded using clusters determined by *Z* > 2.3 and a corrected cluster significance of *p* < 0.05. Locations of slices are indicated by the x, y, and z coordinates as per the MNI coordinate system. (a) Brain regions where the BOLD signal difference between lying and truth telling conditions correlated with at least one of the behavioral indices. Average speed in answering questions truthfully [RT_truth_] was used as representative of participants’ baseline speed. *Relative appended lie RT* [(RT_lie_ − RT_truth_) / RT_truth_] was used as representative of RT-cost of lying. This measure is an indicator of how participants’ RTs changed while lying. **(b)** Brain regions that showed significant BOLD signal difference between lying and truth telling conditions but did not correlate with either of behavioral RT indices. Color bar indicates *Z*-values. ACC, Anterior cingulate cortex; IFG, Inferior frontal gyrus; L, Left; LOC, Lateral occipital cortex; PCC, Posterior cingulate cortex; PCG, Postcentral gyrus; pMTG, Posterior middle temporal gyrus; SMA, Supplementary motor area.

## 4. DISCUSSION

We investigated reaction times and fMRI brain activity of subjects while they provided truthful or intentional false responses to a set of autobiographical yes-no questions. In agreement with multiple previous findings of longer RTs under deceptive conditions (Mameli et al., 2010; Marchewka et al., 2012; Nuñez et al., 2005) our results show that it takes longer to provide a false response than to answer truthfully. However, the size of this effect varied considerably from one participant to another: RT-cost of lying exceeded half a second in some individuals while being absent, and even negative, in some others (Figure 1.b). Inter-individual variability was also present in the baseline speed of participants: the slowest participant required almost triple the time spent by the fastest participant to answer questions truthfully (Figure 1.a). Results of our fMRI comparison between lie and truth conditions (Table 1) resembled previous neuroimaging findings (Christ et al., 2009; Farah et al., 2014), indicating lie-related BOLD signal change in regions of lateral and medial frontal cortex, as well as cingulate, parietal, and insular cortex. We classified this set of regions according to their correlation with behavioral RT-measures for baseline speed and lying RT-cost. Whether a region’s BOLD signal-change during lying associates with RT-prolongation can hint at the probable cognitive function of that region during deception. In what follows, we will discuss general implications of current results in the light of extant literature.

### 4.1. Correlation with RT-cost measure

Our results showed that the amount of increase in BOLD activity of left VLPFC, bilateral ACC, and left LOC areas during lying directly correlates with lying RT-cost measure of each participant. Bilateral PCC and right insula showed inverse correlation with this measure. Correlation between activity of a region and RT-cost measure implies that the cognitive function undertaken by such region is critical for determining the reaction time length in events that required intentional false responding. We standardized RT-costs by expressing the lie-truth RT difference as a fraction of truthful responding speed of each participant. Therefore, if a brain region correlates with RT-cost measure but does not correlate with baseline speed measure the cognitive function of such region is probably exclusively employed for providing intentional false responses, but not truthful answers. Overlap between RT-cost and baseline-speed correlating regions was only observed at the junction of ACC and paracingulate cortex. Pinpointing neural correlates of RT prolongation during lying in prefrontal and executive control regions corroborates the accumulating evidence indicating a predominant role for these regions in deceptive behavior (Abe, 2011; Christ et al., 2009).

Our observation that neural correlates of RT prolongation during lying consists of both negatively correlating and positively correlating areas offers an explanation for the inconsistency of results achieved through transcranial direct current stimulation (tDCS) studies of deception. In first of these studies, Priori et al. (2008) applied tDCS to dorsolateral prefrontal cortex in order to manipulate the excitability of brain regions involved in deception. They found amplified RT-cost of lying after anodal tDCS but no change after cathodal tDCS. In contrast, Karim et al. (2010) found facilitation in lying after cathodal stimulation of anterior prefrontal cortex, as evidenced by reduced RT-cost and lower skin conductance responses, yet no effect after application of anodal tDCS to the same region. Reduction in RT-cost of lying after tDCS is also reported by Mameli et al. (2010) even though they applied anodal tDCS to dorsolateral prefrontal cortex. Our results suggest that slight spatial shifts—in range of centimeters—in the focus of functional changes induced by tDCS can lead to disparate behavioral outcomes. For instance facilitating the excitability of ACC—a region implied in conflict monitoring (Botvinick, Cohen, & Carter, 2004)—could increase the lie RT while the same modulation applied on PCC—a region implied in internally-directed cognition and autobiographical memory retrieval (Leech & Sharp, 2014)—could contrarily decrease the lie RT: since activity within these adjoining regions correlate with RT-cost of lying in opposite directions.

### 4.2. Correlation with baseline-RT measure

In order to represent the dexterity of individuals in performing task instructions, we used average RT of participants during truthful answering as the measure of baseline speed. Participants who had faster baseline speed showed larger activity increase in right VLPFC and bilateral paracingulate cortex during lying. Correlation with baseline-RT implies involvement with cognitive functions that are shared between truth telling and lying events, but employed to a larger extent during lying. This implication of our results conforms to previous literature suggesting general, nonspecific executive function roles for right VLPFC and paracingulate cortex. A recent meta-analysis (Levy & Wagner, 2011) found functional specialization within right VLPFC for detection of behaviorally relevant stimuli, updating of action plans, and responding to decision uncertainty: functions that are employed during a wide variety of cognitive tasks including truth telling and lying. More in line with the inter-individual difference approach of our study, Fornito et al. (2004) has shown that prominent paracingulate sulcus folding in an individual is associated with non-specific performance advantage on cognitively demanding executive function tasks. Our results suggest that paracingulate and right VLPFC functions relate to truth and lie RTs similarly and cannot impose a lie RT-cost in the same way that lie exclusive cognitive processes do.

### 4.3. Lie-related regions not correlating with either RT measures

In bilateral SMA, ICWM and left AG, lie-related BOLD increase did not correlate with either of behavioral RT measures. This implies that the function undertaken by these areas does not contribute to prolongation of reaction times. These regions might perform relatively fast cognitive processes and/or work in parallel with other processes. Alternatively, such a region might be activated subsequent to subjects’ response and play roles in retrospective evaluation of action. It is notable that both cortical areas in this subset associate with higher-level motor functions: subregions within SMA are involved with multiple stages of movement from preparation to execution (K. M. Lee, Chang, & Roh, 1999) and AG is believed to represent action awareness (Farrer et al., 2008). Our paradigm used two-alternative choice questions. It is conceivable that participants might have employed a motor task-switching strategy for providing false responses. Such strategy could justify higher activation in motor control regions in parallel and subsequent to cognitive processes culminating in execution of a deceptive behavior.

### 4.4. Functional dichotomy in VLPFC and cingulate cortex

A noteworthy finding of our study is the dichotomy between right and left VLPFC, which accordingly showed exclusive correlation with baseline-speed and lying RT-cost measures. Our findings imply that the cognitive function of right VLPFC is shared between lie and truth events but the function of left VLPFC is more specific to lying. This is consistent with previous literature signifying the functional dissociation between contralateral VLPFCs: right VLPFC is assumed to respond to decision uncertainty and motor inhibition (Levy & Wagner, 2011), while data from left VLPFC support a role in cognitive control of memory (Badre & Wagner, 2007). Areas in medial frontal and cingulate cortex also revealed an interesting pattern of functional dissociation. From anterior to posterior, ACC correlated with lie RT-cost positively, paracingulate region correlated with baseline speed, and PCC correlated with RT-cost negatively. This finding is in line with the alleged fundamental dichotomy within cingulate cortex, with anterior executive and posterior evaluative functions (Mohanty et al., 2007; Vogt, Finch, & Olson, 1992).

### 4.5. Time-on-task effects might contaminate fMRI comparisons between lie and truth

It should be noted that fMRI results we discussed so far are correlative by nature; therefore, the direction of causality in the observed correlations could not be readily inferred. So far, we strived to interpret these correlations as the neural correlates of reaction time prolongation during lying; however, we should also consider the reverse causation direction: the possibility that activity of a brain region be modulated as a consequence of longer reaction times during lying. In fMRI experiments, participants briskly employ resources to responds to an event but are free to disengage in rest periods interleaving trials. A longer reaction time, whatever be the mechanism underlying its prolongation, calls for higher amount of maintained attention and goal-directed behavior (Grinband, Wager, Lindquist, Ferrera, & Hirsch, 2008). Indeed, recent studies have found that the length of an individual’s “time-on-task” monotonically increases the BOLD amplitude within multiple frontal and parietal regions, irrespective of the nature of cognitive task at hand (Grinband et al., 2011; Yarkoni, Barch, Gray, Conturo, & Braver, 2009). The extent of resemblance between brain regions affected by time-on-task duration (Yarkoni et al., 2009) and regions consistently reported by fMRI studies of deception (Farah et al., 2014) is remarkable. Bilateral medial frontal gyrus, right IFG/VLPFC, right middle frontal gyrus/DLPFC, left anterior insula, left IPL, left precuneus/intraparietal sulcus, and bilateral thalamus show up in both lists. Nevertheless, ACC, PCC, and left VLPFC are among the deception-related regions that have not been implicated in time-on-task effects. In the current study, we found a similar dichotomy by separating deception-related regions correlating with the baseline speed of participants from regions correlating with the lying RT-cost. The fact that only right VLPFC is reported to show time-on-task effect bolsters the conjecture of left-lateralized involvement of VLPFC in deception that we proposed above.

On a wider perspective, the supposition that time-on-task duration affects frontal activity can undermine the validity of current mapping of the neural correlates of deception. Most of our knowledge comes from fMRI studies contrasting brain activity during lying versus truthful responding, without controlling for the confounding factor of difference in reaction times under the two conditions. By reviewing the studies included in the latest meta-analysis (Farah et al., 2014) we saw that only two out of 23 studies have included reaction times in their fMRI model (Browndyke et al., 2008; Nuñez et al., 2005) despite the fact that most studies did report significantly longer RTs under deception conditions (see Supplementary information 1). Due to this systematic shortcoming, set of brain regions currently associated with deceptive behavior is probably contaminated by areas that show time-on-task effects but do not necessarily play a critical part in generation of deception. Further studies should address this ambiguity by following a more controlled approach to brain mapping of deception.

### 4.6. Limitations and future suggestions

We acknowledge that the two-alternative forced choice questions used in this study does not tap on the whole spectrum of processes involved in deception. On events requiring intentional false responding, participants probably relied largely on response switching strategies: swapping the truthful response with the false alternative at the last moment. Mental processes required for fabrication of deceptive responses and cognitive planning are in all likelihood not engaged during the course of our experiment. Emotional engagement that is normally experienced during real deceptive acts was also probably absent in the current study: our subjects were asked to blindly select half of questions and were later instructed to lie about them, while in ecologically valid situations the decision to lie will be determined by hidden personal motives and incentives. In addition to the type of questions, the verbal nature of them should be noted too. Our results implied left-lateralized specification of VLPFC for deception; however, it is unclear whether left-lateralization would be replicated in case of non-verbal forms of deception.

In the current discussion we tried to propose provisional implications regarding the probable cognitive function of lie-related brain areas based on the pattern of correlation with RT measures; nevertheless, we should reiterate that our experiment was not designed to provide exact inference about cognitive functions. Further studies are called for to confirm current implications. In this study, we exploited the between-subject variability in reaction times; a future study can try to investigate the within-subject trial-by-trial variability of reaction times and their correlation with activity level of lie-related brain regions.

### 4.7. Conclusion

In this study, we tried to find what neural components contribute to RT prolongation when people try to respond deceptively. Based on current results, we speculate that cognitive functions undertaken by left VLPFC and cingulate cortex—regions correlating with the RT-cost measure—determine the amount of RT prolongation during lying and therefore might be more critical for producing deception. On the other hand, the increase in activation of paracingulate and right VLPFC—areas that correlated with the baseline speed measure and implicated in the time-on-task effect by the literature—might be mere byproducts of longer reaction times and higher mental load during deception.

## 5. ACKNOWLEDGEMENTS

Authors report no conflicts of interest. This research was supported by a grant from Tehran University of Medical Sciences.

## REFERENCES

Abe, N. (2011). How the Brain Shapes Deception: An Integrated Review of the Literature. The Neuroscientist, 17(5), 560–574. http://doi.org/10.1177/1073858410393359

Badre, D., & Wagner, A. D. (2007). Left ventrolateral prefrontal cortex and the cognitive control of memory. Neuropsychologia, 45(13), 2883–901. http://doi.org/10.1016/j.neuropsychologia.2007.06.015

Botvinick, M. M., Cohen, J. D., & Carter, C. S. (2004). Conflict monitoring and anterior cingulate cortex: An update. Trends in Cognitive Sciences. http://doi.org/10.1016/j.tics.2004.10.003

Browndyke, J. N., Paskavitz, J., Sweet, L. H., Cohen, R. A., Tucker, K. A., Welsh-Bohmer, K. A.,… Schmechel, D. E. (2008). Neuroanatomical correlates of malingered memory impairment: event-related fMRI of deception on a recognition memory task. Brain Injury, 22(6), 481–9. http://doi.org/10.1080/02699050802084894

Caso, L., Gnisci, A., Vrij, A., & Mann, S. (2005). Processes underlying deception: an empirical analysis of truth and lies when manipulating the stakes. Journal of Investigative…, 2(3), 195–202. http://doi.org/10.1002/jip.32

Christ, S. E., Van Essen, D. C., Watson, J. M., Brubaker, L. E., & McDermott, K. B. (2009). The contributions of prefrontal cortex and executive control to deception: Evidence from activation likelihood estimate meta-analyses. Cerebral Cortex, 19(7), 1557–1566. http://doi.org/10.1093/cercor/bhn189

Debey, E., De Houwer, J., & Verschuere, B. (2014). Lying relies on the truth. Cognition, 132(3), 324–334. http://doi.org/10.1016/j.cognition.2014.04.009

Debey, E., Liefooghe, B., De Houwer, J., & Verschuere, B. (2014). Lie, truth, lie: the role of task switching in a deception context. Psychological Research, 478–488. http://doi.org/10.1007/s00426-014-0582-4

Farah, M. J., Hutchinson, J. B., Phelps, E. a, & Wagner, A. D. (2014). Functional MRI-based lie detection: scientific and societal challenges. Nature Publishing Group, 15(2), 123–131. http://doi.org/10.1038/nrn3665

Farrer, C., Frey, S. H., Van Horn, J. D., Tunik, E., Turk, D., Inati, S., & Grafton, S. T. (2008). The angular gyrus computes action awareness representations. Cerebral Cortex, 18(2), 254–261. http://doi.org/10.1093/cercor/bhm050

Farrow, T. F. D., Hopwood, M.-C. C., Parks, R. W., Hunter, M. D., & Spence, S. A. (2011). Evidence of mnemonic ability selectively affecting truthful and deceptive response dynamics. American Journal of Psychology, 124(4), 447–453. http://doi.org/10.5406/amerjpsyc.124.4.0447

Fornito, A., Yucel, M., Wood, S., Stuart, G. W., Buchanan, J. A., Proffitt, T.,… Pantelis, C. (2004). Individual Differences in Anterior Cingulate/Paracingulate Morphology Are Related to Executive Functions in Healthy Males. Cerebral Cortex, 14(4), 424–431. http://doi.org/10.1093/cercor/bhh004

Ganis, G., Kosslyn, S. M., Stose, S., Thompson, W. L., & Yurgelun-Todd, D. A. (2003). Neural correlates of different types of deception: An fMRI investigation. Cerebral Cortex, 13(8), 830–836. http://doi.org/10.1093/cercor/13.8.830

Gombos, V. A. V. A. (2006). The Cognition of Deception: The Role of Executive Processes in Producing Lies. Genetic, Social and General Psychology Monographs, 132(3), 197–214. http://doi.org/10.3200/MONO.132.3.197-214

Grinband, J., Savitskaya, J., Wager, T. D., Teichert, T., Ferrera, V. P., & Hirsch, J. (2011). The dorsal medial frontal cortex is sensitive to time on task, not response conflict or error likelihood. NeuroImage, 57(2), 303–311. http://doi.org/10.1016/j.neuroimage.2010.12.027

Grinband, J., Wager, T. D., Lindquist, M., Ferrera, V. P., & Hirsch, J. (2008). Detection of time-varying signals in event-related fMRI designs. NeuroImage, 43(3), 509–520. http://doi.org/10.1016/j.neuroimage.2008.07.065

Jenkinson, M., Bannister, P., Brady, M., & Smith, S. (2002). Improved Optimization for the Robust and Accurate Linear Registration and Motion Correction of Brain Images. NeuroImage, 17(2), 825–841. http://doi.org/10.1006/nimg.2002.1132

Karim, A. A., Schneider, M., Lotze, M., Veit, R., Sauseng, P., Braun, C., & Birbaumer, N. (2010). The Truth about Lying: Inhibition of the Anterior Prefrontal Cortex Improves Deceptive Behavior. Cerebral Cortex, 20(1), 205–213. http://doi.org/10.1093/cercor/bhp090

Kleinberg, B., & Verschuere, B. (2016). The role of motivation to avoid detection in reaction time-based concealed information detection. Journal of Applied Research in Memory and Cognition, 5(1), 43–51. http://doi.org/10.1016/j.jarmac.2015.11.004

Kozel, F. A., Johnson, K. A., Grenesko, E. L., Laken, S. J., Kose, S., Lu, X.,… George, M. S. (2009). Functional MRI detection of deception after committing a mock sabotage crime. In Journal of Forensic Sciences (Vol. 54, pp. 220–231). http://doi.org/10.1111/j.1556-4029.2008.00927.x

Kozel, F. A., Johnson, K. A., Mu, Q., Grenesko, E. L., Laken, S. J., & George, M. S. (2005). Detecting deception using functional magnetic resonance imaging. Biological Psychiatry, 58(8), 605–13. http://doi.org/10.1016/j.biopsych.2005.07.040

Langleben, D. D., Schroeder, L., Maldjian, J. A., Gur, R. C., McDonald, S., Ragland, J. D.,… Childress, A. R. (2002). Brain activity during simulated deception: an event-related functional magnetic resonance study. NeuroImage, 15(3), 727–32. http://doi.org/10.1006/nimg.2001.1003

Lee, K. M., Chang, K. H., & Roh, J. K. (1999). Subregions within the supplementary motor area activated at different stages of movement preparation and execution. NeuroImage, 9(1), 117–123. http://doi.org/10.1006/nimg.1998.0393

Lee, T. M. C., Liu, H.-L., Tan, L.-H., Chan, C. C. H., Mahankali, S., Feng, C.-M.,… Gao, J.-H. (2002). Lie detection by functional magnetic resonance imaging. Human Brain Mapping, 15(3), 157–164. http://doi.org/10.1002/hbm.10020

Leech, R., & Sharp, D. J. (2014). The role of the posterior cingulate cortex in cognition and disease. Brain. http://doi.org/10.1093/brain/awt162

Levy, B. J., & Wagner, A. D. (2011). Cognitive control and right ventrolateral prefrontal cortex: reflexive reorienting, motor inhibition, and action updating. Annals of the New York Academy of Sciences, 1224(Ba 45), 40–62. http://doi.org/10.1111/j.1749-6632.2011.05958.x

Mameli, F., Mrakic-Sposta, S., Vergari, M., Fumagalli, M., Macis, M., Ferrucci, R.,… Priori, A. (2010). Dorsolateral prefrontal cortex specifically processes general - but not personal - knowledge deception: Multiple brain networks for lying. Behavioural Brain Research, 211(2), 164–168. http://doi.org/10.1016/j.bbr.2010.03.024

Marchewka, A., Jednorog, K., Falkiewicz, M., Szeszkowski, W., Grabowska, A., & Szatkowska, I. (2012). Sex, lies and fMRI--gender differences in neural basis of deception. PloS One, 7(8), e43076. http://doi.org/10.1371/journal.pone.0043076

Marston, W. M. (1920). Reaction-time symptoms of deception. Journal of Experimental Psychology, 3(1), 72–87. http://doi.org/10.1037/h0067963

Mohanty, A., Engels, A. S., Herrington, J. D., Heller, W., Ho, M.-H. R., Banich, M. T.,… Miller, G. a. (2007). Differential engagement of anterior cingulate cortex subdivisions for cognitive and emotional function. Psychophysiology, 44(3), 343–51. http://doi.org/10.1111/j.1469-8986.2007.00515.x

Mumford, J. A., Poline, J.-B., & Poldrack, R. A. (2015). Orthogonalization of Regressors in fMRI Models. PLOS ONE, 10(4), e0126255. http://doi.org/10.1371/journal.pone.0126255

Nuñez, J. M., Casey, B. J., Egner, T., Hare, T., & Hirsch, J. (2005). Intentional false responding shares neural substrates with response conflict and cognitive control. NeuroImage, 25(1), 267–77. http://doi.org/10.1016/j.neuroimage.2004.10.041

Priori, A., Mameli, F., Cogiamanian, F., Marceglia, S., Tiriticco, M., Mrakic-Sposta, S.,… Sartori, G. (2008). Lie-specific involvement of dorsolateral prefrontal cortex in deception. Cerebral Cortex, 18(2), 451–455. http://doi.org/10.1093/cercor/bhm088

Sheridan, M. R., & Flowers, K. A. (2010). Reaction times and deception: The lying constant. International Journal of Psycholgical Studies, 2(2), 41–51. http://doi.org/10.5539/ijps.v2n2p41

Sip, K. E., Lynge, M., Wallentin, M., McGregor, W. B., Frith, C. D., & Roepstorff, A. (2010). The production and detection of deception in an interactive game. Neuropsychologia, 48(12), 3619–3626. http://doi.org/10.1016/j.neuropsychologia.2010.08.013

Sip, K. E., Roepstorff, A., McGregor, W., & Frith, C. D. (2008). Detecting deception: the scope and limits. Trends in Cognitive Sciences, 12(2), 48–53. http://doi.org/10.1016/j.tics.2007.11.008

Sip, K. E., Skewes, J. C., Marchant, J. L., McGregor, W. B., Roepstorff, A., & Frith, C. D. (2012). What if I get busted? Deception, choice, and decision-making in social interaction. Frontiers in Neuroscience, (APR). http://doi.org/10.3389/fnins.2012.00058

Smith, S. M. (2002). Fast robust automated brain extraction. Human Brain Mapping, 17(3), 143–55. http://doi.org/10.1002/hbm.10062

Spence, S. A., Farrow, T. F. D., Herford, A. E., Wilkinson, I. D., Zheng, Y., & Woodruff, P. W. R. (2001). Behavioural and functional anatomical correlates of deception in humans. Neuroreport, 12(13), 2849–2853. http://doi.org/10.1097/00001756-200109170-00019

Varga, M., Visu-Petra, G., Miclea, M., & Visu-Petra, L. (2015). The “good cop, bad cop” effect in the rt-based concealed information test: Exploring the effect of emotional expressions displayed by a virtual investigator. PLoS ONE, 10(2). http://doi.org/10.1371/journal.pone.0116087

Vendemia, J. M. C., Buzan, R. F., & Simon-Dack, S. L. (2005). Reaction time of motor responses in two-stimulus paradigms involving deception and congruity with varying levels of difficulty. Behavioural Neurology, 16, 25–36. http://doi.org/10.1155/2005/804026

Verschuere, B., & in ´t Hout, W. (2016). Psychopathic Traits and Their Relationship with the Cognitive Costs and Compulsive Nature of Lying in Offenders. PLOS ONE, 11(7), e0158595. http://doi.org/10.1371/journal.pone.0158595

Verschuere, B., Suchotzki, K., & Debey, E. (2015). Detecting Deception Through Reaction Times. In Detecting Deception: Current Challenges and Cognitive Approaches (pp. 269–291). http://doi.org/10.1002/9781118510001.ch12

Visu-Petra, G., Miclea, M., Buş, I., & Visu-Petra, L. (2014). Detecting concealed information: The role of individual differences in executive functions and social desirability. Psychology, Crime & Law, 20(1), 20–36. http://doi.org/10.1080/1068316X.2012.736509

Visu-Petra, G., Miclea, M., & Visu-Petra, L. (2012). Reaction Time-based Detection of Concealed Information in Relation to Individual Differences in Executive Functioning. Applied Cognitive Psychology, 26(3), 342–351. http://doi.org/10.1002/acp.1827

Vogt, B. a, Finch, D. M., & Olson, C. R. (1992). Functional heterogeneity in cingulate cortex: the anterior executive and posterior evaluative regions. Cerebral Cortex (New York, N.Y.□: 1991), 2(6), 435–43. Retrieved from http://www.ncbi.nlm.nih.gov/pubmed/1477524

Vrij, A., Granhag, P. A., Mann, S., & Leal, S. (2011). Outsmarting the Liars: Toward a Cognitive Lie Detection Approach. Current Directions in…, 20(1), 28–32. http://doi.org/10.1177/0963721410391245

Walczyk, J. J., Roper, K. S., Seemann, E., & Humphrey, A. M. (2003). Cognitive mechanisms underlying lying to questions: response time as a cue to deception. Applied Cognitive Psychology, 17(7), 755–774. http://doi.org/10.1002/acp.914

Williams, E. J., Bott, L. A., Patrick, J., & Lewis, M. B. (2013). Telling Lies: The Irrepressible Truth? PLoS ONE, 8(4). http://doi.org/10.1371/journal.pone.0060713

Woolrich, M. W., Behrens, T. E. J., Beckmann, C. F., Jenkinson, M., & Smith, S. M. (2004). Multilevel linear modelling for FMRI group analysis using Bayesian inference. NeuroImage, 21(4), 1732–1747.

Yarkoni, T., Barch, D. M., Gray, J. R., Conturo, T. E., & Braver, T. S. (2009). BOLD Correlates of Trial-by-Trial Reaction Time Variability in Gray and White Matter: A Multi-Study fMRI Analysis. PLoS ONE, 4(1), e4257. http://doi.org/10.1371/journal.pone.0004257

